# The mitochondrial genome segregation system of *T. brucei*: Single p197 molecules connect the basal body with the outer membrane

**DOI:** 10.1101/2022.03.10.483810

**Authors:** Salome Aeschlimann, Ana Kalichava, Bernd Schimanski, Philip Stettler, Torsten Ochsenreiter, André Schneider

**Affiliations:** Department of Chemistry, Biochemistry and Pharmaceutical Sciences, University of Bern, Freiestrasse 3, Bern CH-3012, Switzerland; Graduate School for Cellular and Biomedical Sciences, University of Bern, Bern, Switzerland; Institute of Cell Biology, University of Bern, Bern CH-3012, Switzerland

## Abstract

The tripartite attachment complex (TAC) couples the segregation of the single unit mitochondrial DNA of trypanosomes with the basal body of the flagellum. Here we studied the architecture of the exclusion zone filament of the TAC that connects the basal body with the mitochondrial outer membrane. The only known component of the exclusion zone filaments is p197. Using genetical, biochemical and microscopical methods we show that p197 has three domains all of which are essential for mitochondrial DNA inheritance. The C-terminus of p197 interacts with the mature and pro-basal body whereas its N-terminus binds to the peripheral outer membrane protein TAC65. The large central region of p197 has a high α-helical content and likely acts as a flexible spacer. Replacement of endogenous p197 with a functional version containing N- and C-terminal epitope tags together with expansion microscopy demonstrates that p197 alone can bridge the approximately 170 nm gap between the basal body and the periphery of the outer membrane. This demonstrates the power of expansion microscopy which allows to localize distinct regions within the same molecule and suggests that p197 is the TAC subunit most proximal to the basal body.

**Significance statement:** Segregation of the replicated single unit mitochondrial genome of *Trypanosoma brucei* requires a large hardwired structure that connects the organellar DNA with the flagellar basal body. The cytosolic part of this structure consists of filaments made of single p197 molecules, a protein larger than 600 kDa. p197 has three domains all of which are essential for its function. The N-terminus of p197 is anchored to the peripheral outer membrane protein TAC65 whereas its C-terminus connects to the base of the basal body. The large central domain forms an α-helix and consists of at least 26 repeats of 175 aa in length. It provides a flexible linker bridging the approximately 170 nm between the outer membrane and the basal body

## Introduction

Replication and segregation of the mitochondrial DNA is essential for proper organellar function. While these processes have been studied in detail in the phylogenetically related yeast and mammals (1, 2), which include most of the popular model systems for cell biology, we know much less about them in protozoans which make up most of the eukaryotic diversity.

An especially interesting case is the parasitic protozoan *Trypanosoma brucei*. Unlike most other eukaryotes it has a single mitochondrion which contains a single unit mitochondrial genome, termed kinetoplastid DNA (kDNA) (3, 4). The kDNA consists of a few dozen maxicircles of 22 kb in length and several thousand minicircles of 1 kb in length which are heterogeneous in sequence. The maxicircle encodes 18 protein coding genes, two rRNAs and corresponds to the mitochondrial DNA found in other species (5, 6). However, many of these genes are cryptogenes meaning that their transcripts need to be edited by multiple uridine insertions and/or deletions to become functional mRNAs (7–9). Maxicircle and minicircle are topologically highly interlocked forming the disc-shaped kDNA network that is localized to a specific region within the mitochondrion opposite of the basal body (BB) of the flagellum.

As a consequence of its single unit nature, and in contrast to the mitochondrial genomes of most other eukaryotes, the kDNA is replicated at a defined time point during the cell cycle just before the onset of the nuclear S-phase (10, 11). Moreover, trypanosomes require precise mechanisms of replication and subsequent segregation of the kDNA which guarantee, that after binary fission of the mitochondrion (12), the two daughters receive one copy each of the replicated kDNA discs in their organelles. Trypanosomes achieve this by coupling kDNA segregation to the segregation of another single copy structure in trypanosomes, the BB of the flagellum (13). This coupling requires a unique structure termed tripartite attachment complex (TAC) which physically connects the kDNA disc, across the two mitochondrial membranes, with the BB of the flagellum. Disruption of the TAC is lethal. It leads to overreplicated kDNA that cannot be segregated anymore eventually resulting in daughter cells lacking kDNA (6, 14).

The TAC is a complicated structure that has three morphologically defined subdomains: (i) the unilateral filaments (ULF) that connect the kDNA to the mitochondrial inner membrane (IM) which is approximately 100 nm in length, (ii) the exclusion zone filaments (EZFs) between the BB of the flagellum and the mitochondrial outer membrane (OM) bridging a distance of about 150-170 nm, and (iii) the differentiated membranes (DMs) that link the ULFs across the two membranes with the EZFs. The DM region is very thin and consists essentially only of the OM and the IM (15). While the TAC is specific for kinetoplastids, recent work suggests that the concept of coupling inheritance of organelles to flagellar segregation also applies to mitosomes of the anaerobic protist *Giardia*. In these cells a subgroup of mitosomes is linked to the axonemes of the oldest flagella (16, 17). This indicates that a structure of unknown composition that is analogous to the trypanosomal EZFs and that links the flagella to the outer membrane of the mitosomes, must exist in *Giardia*. In recent years much progress has been made identifying the components of the different TAC subdomains. The ULF region was shown to consist of p166, the first TAC component identified (18) which also connects to the IM (19), and of TAC102 which localizes close to the kDNA (20). Counterintuitively, the DMs, the smallest subregion of the TAC, has the most complex composition. In the OM alone four different integral membrane proteins (TAC60, TAC40, TAC42 and pATOM36) as well as TAC65, a peripherally OM-associated protein that faces the cytosol, have been identified. All of these TAC subunits are essential for TAC function (21, 22). Moreover, pATOM36 in addition to its role in the TAC is also involved in the biogenesis of OM proteins (23). It is a functional analogue of the yeast MIM complex (24) and thus integrates mitochondrial protein import with kDNA inheritance (25). In contrast, p166 is the only TAC subunit that has been characterized in the mitochondrial IM (19). The EZF subdomain of the TAC finally consists of at least one essential factor: p197 which is the focus of the present study (26).

The TAC is unusual compared to other mitochondrial protein complexes as there is only a single such structure in the mitochondrion of non-dividing cells (15). This raises the question of how it is formed? A recent analysis revealed that the overarching principle of TAC biogenesis is a hierarchical assembly starting from the BB of the flagellum progressing towards the kDNA (27).

A prerequisite to build a precise model of the TAC and to understand how the structure is assembled requires information about how each TAC component is arranged within the structure and how it interacts with which other TAC components. Here we provide this information for p197, the largest TAC component. According to TriTrypDB p197 has a predicted molecular mass of 197 kDa and contains 3.5 tandem repeats of 175 identical aa. Using molecular genetics and in vivo complementation assays, involving differentially tagged p197 variants combined with ultrastructure expansion microscopy (U-ExM), we show that assembly of p197 and thus of the whole TAC starts very early in the cell cycle immediately after the formation of the pro-basal body (pro-BB). We define which domains of p197 are essential for localization and/or function of p197. We demonstrate that individual molecules of p197 form the EZF. Finally, we show how p197 is arranged in this TAC region and identify its interaction partner in the periphery of the OM.

## Results

### p197-HA is functional and localizes to both mature and pro-BBs

To investigate the role of p197 in the TAC, we produced an RNAi cell line targeting the 3’UTR of p197 (p197 3’UTR RNAi). In this cell line induction of RNAi strongly interferes with normal growth and causes mis-segregation of the kDNA resulting in cells that either lack kDNA or contain overreplicated kDNA networks (*SI Appendix*, Fig. S1A). This is in agreement with previous studies (26) and corresponds to the typical phenotype of cell lines that are deficient for TAC subunits (14). Since RNAi targets the 3’UTR of the p197 mRNA its ORF is not affected. Thus, we used the p197 3’UTR RNAi cell line and in situ HA-tagged one of its p197 alleles at the C-terminus. (Note that the tagging replaces the endogenous 3’UTR). Addition of tetracycline (Tet) to this cell line, termed p197 3’UTR RNAi + p197-HA, shows that expression of the RNAi-resistant p197-HA version restores normal growth and kDNA segregation indicating that the C-terminal HA tag does not interfere with p197 function (*SI Appendix*, Fig. S1B).

Subsequently, immunofluorescence (IF) analysis of detergent-extracted cytoskeletons was used to compare the localization of p197-HA with the trypanosomal SAS-6 homologue (Fig. 1A). SAS-6 is an evolutionary conserved protein and has been implicated in the assembly of the centriole cartwheel (28). In *T. brucei* SAS-6 localizes to both the mature BB as well as the pro-BB throughout the cell cycle (29–31).

**Figure 1.**
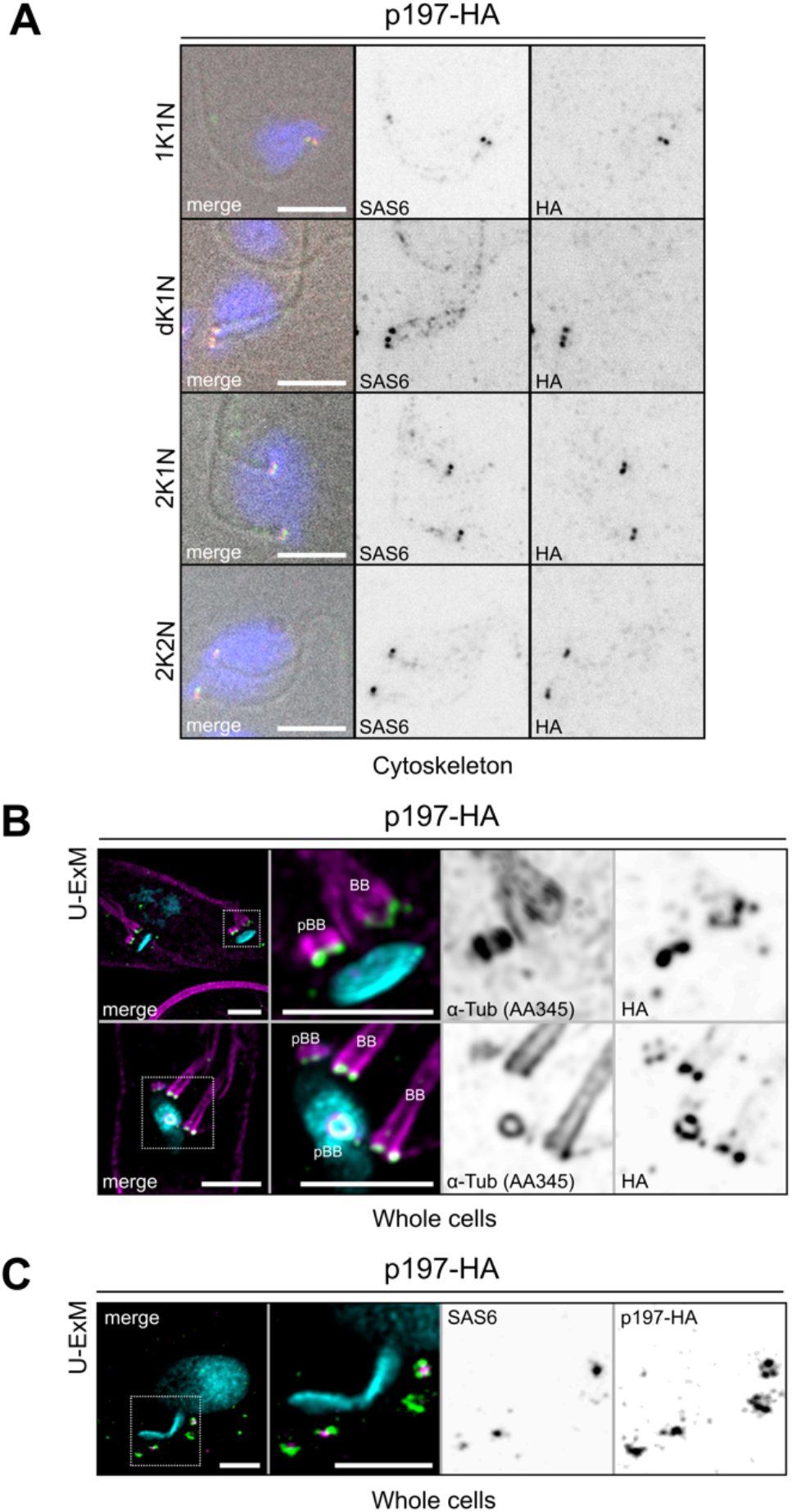
p197 localizes to both the BB and pro-BB throughout the cell cycle. **(A)** IF analysis of cytoskeletons isolated from a p197-HA expressing cell stained with DAPI (blue), for the BB and pro-BB marker SAS6 (green) and for HA (red) as indicated. Representative images of the cell cycle stages with kDNA (K) nucleus (N) configurations of 1K1N, dK1N, 2K1N and 2K2N are shown. Scale bar 5μm **(B)** U-ExM of p197-HA expressing cells stained with DAPI, the recombinant α-tubulin antibody AA345 (magenta) and for HA (green) as indicated. The upper row shows a 2K1N and the lower one a dK1N cell cycle stage. Scale bar 2μm. **(C)** U-ExM as in (B) but cells were stained with DAPI (blue), for SAS6 (magenta) and HA (green). Scale bar 2μm.

In interphase cells, with one kDNA (K) and one Nucleus (N), a configuration termed 1K1N, both proteins colocalize to two closely opposed dots that correspond to the mature BB and the pro-BB (Fig. 1A, top row), respectively. Further into the cell cycle, when the kDNA has been replicated but not yet segregated (configuration: dK1N), the two proteins localize to three adjacent dots in most cells, likely corresponding to the mature BB and pro-BB of the old flagellum and the new mature BB of the new flagellum (Fig. 1A, second row). Finally, two dots each of p197 and SAS-6 are detected for each kDNA in cells showing 2K1N and 2K2N configurations (Fig. 1A, last two rows). Thus, p197-HA colocalizes with SAS-6 throughout the cell cycle, indicating that assembly of the TAC starts simultaneously with the formation of the pro-BB and thus precedes the growth of the new flagellum as has been reported recently (32).

### p197 localizes to the bottom of the BB surrounding SAS-6

To more precisely analyze the intracellular localization of p197-HA we used U-ExM which was recently established for trypanosomes (33–35). U-ExM allows to increase the spatial resolution for microscopic imaging by physical expansion of the cell (36, 37). Figure 1B shows the cellular region of *T. brucei* that contains the BBs and the kDNAs. It is stained for p197-HA and α-tubulin using the antibody AA345 (38), which detects the mature and pro-BB (27, 39). The resolution of the pictures is greatly increased when compared to conventional IF microscopy. As expected for the mature and the pro-BB the α-tubulin signals are consistent with a hollow cylindrical structure which is viewed either from the top, the bottom, or from the side. p197 is detected adjacent to the bottom layer of the mature BB and the pro-BB and seems to form a partial ring that extends the hollow cylinder defined by the 9 microtubule triplets of the BB. The U-ExM in Figure 1C shows that, in agreement with its proposed function in cartwheel assembly, SAS-6 localizes to the center of the partial ring defined by p197 (33). In summary, these results suggest that p197 is the TAC subunit that is most proximal to the mature BB and the pro-BB.

### The C-terminus of p197 is anchored at the BB

Next we complemented the p197 3’UTR RNAi cell line with a Tet-inducible C-terminal HA-tagged p197 variant lacking the N-terminal 1228 aa (ΔN1228-HA). Figure 2A demonstrates that expression of ΔN1228-HA cannot restore the growth inhibition caused by the absence of full length p197. However, ΔN1228-HA correctly localizes to both the mature and the pro-BBs, stained by the α-tubulin antibody YL1/2 (Fig. 2B), even though the cell lacks kDNA due to the absence of the TAC (Fig. 2B, bottom, panel). Thus, while ΔN1228-HA cannot assemble a functional TAC the C-terminal 520 aa of p197 appear to be sufficient to interact with the mature and the pro-BBs.

**Figure 2.**
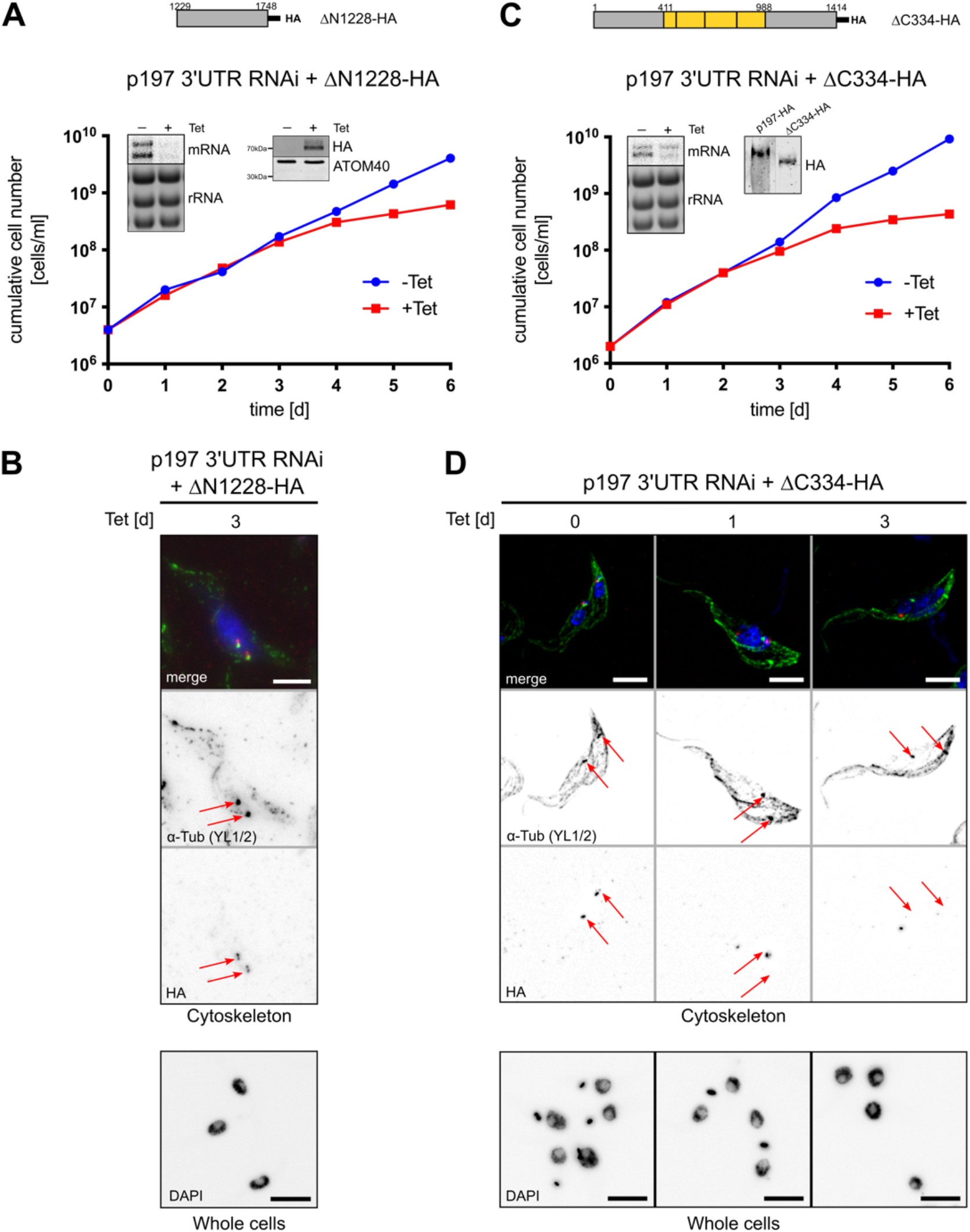
The C-terminus of p197 is anchored at the BB. **(A)** Growth curve of the p197 3’UTR RNAi cell line expressing an RNAi resistant in-situ C-terminally HA tagged p197 version lacking the 1228 N-terminal aa, termed ΔN1228-HA, schematically depicted at the top. Inset left, Northern blot showing ablation of p197 mRNA, ethidiumbromide-stained rRNAs serve as loading control. Inset right, immunoblot showing Tet-inducible expression of ΔN1228-HA. ATOM40 serves as loading control. **(B)** Top panels, IF analysis of cytoskeletons isolated from the cell line shown in (A) stained with DAPI (blue), the α-tubulin antibody YL1/2 (green) and for HA (red) as indicated. Time of RNAi induction is indicated. Red arrows point towards the two basal bodies. Bottom panel, IF analysis of whole cells corresponding to the top panels stained with DAPI to illustrate the RNAi-induced kDNA loss. Scale bar 5μm. **(C)** Growth curve of the p197 3’UTR RNAi cell line expressing an RNAi resistant in situ C-terminally HA tagged p197 version lacking the 334 C-terminal aa, termed ΔC334-HA, schematically depicted at the top. Inset left, Northern blot showing ablation of p197 mRNA, ethidiumbromide-stained rRNAs serve as loading control. Inset right, immunoblot showing constitutive expression of ΔC334-HA. Full-length p197 serves as size marker for comparison. **(D)** Top panels, IF analysis of cytoskeletons isolated from the cell line shown in (C) stained with DAPI (blue), the α-tubulin antibody YL1/2 (green) and for HA (red) as indicated. Time of RNAi induction is indicated. Red arrows point towards the two basal bodies. Bottom panel, IF analysis of whole cells corresponding to the top panels stained with DAPI to illustrate the RNAi-induced kDNA loss. Scale bar 5μm.

To confirm and extend these results we used the p197 3’UTR RNAi cell line and modified one allele of p197 by in situ tagging so that it lacks the C-terminal 334 aa and contains a C-terminal HA-tag (ΔC334-HA). The resulting cell line was termed p197 3’UTR RNA + ΔC334-HA. However, as for the p197 3’UTR RNAi + ΔN1228-HA cell line described above, the ΔC334-HA variant does not restore normal growth and thus cannot functionally replace full length p197 (Fig. 2C). IF analyses show that in uninduced RNAi cells, which contain intact and fully functional TAC structures, ΔC334-HA is correctly localized to the BB region. However, while ΔC334-HA localizes to the preexisting TAC at the old BB, which is still connected to the kDNA, after one day of p197 RNAi induction, it is absent from the newly formed BB, which lacks a TAC and therefore the kDNA (Fig. 2D). After three days of RNAi induction, finally, when the slow growth phenotype becomes apparent and the kDNA and the TAC has been completely lost, the ΔC334-HA, while still detected by SDS PAGE, is absent from the BB region (Fig. 2D). These results show that in the presence of an intact TAC ΔC334-HA is correctly localized. Thus ΔC334-HA has either retained its capability to connect to its downstream binding partner that mediates the interaction of p197 with the OM or it interacts laterally with full length p197 present in the old TAC.

In summary these results indicate that the C-terminal 334 to 520 aa of p197 are necessary and sufficient for the upstream interaction of p197 with the BBs.

### The N-terminus of p197 is anchored at the OM

To investigate the two possibilities raised above, we generated a more drastic C-terminal deletion mutant of p197 in which 1383 aa of its C-terminus were deleted. Similar to the ΔC334-HA variant the ΔC1383-HA variant is not functional yet able to correctly localize in cells that still have full length p197. (Fig. 3A, B). We therefore expressed the ΔC1383-HA variant in a cell line capable of inducible RNAi of the OM TAC subunit TAC40 (Fig. 3C). Flagella and the TAC are stable structures that remain connected to each other during a standard flagellar isolation procedure (13, 21). Fig. 3C shows that in the uninduced TAC40 RNAi cell line ΔC1383-HA was detected in the BB region in 80% of isolated flagella, indicating that as shown above (Fig. 3A, B) non functional ΔC1383-HA still localizes to the TAC. Ablation of TAC40 disrupts the formation of the TAC part downstream of TAC40 resulting in the delocalization of TAC components, such as p166 and TAC102 (27). Moreover, also the OM TAC subunit TAC60 and the cytosolic TAC65 that is peripherally attached to the OM get delocalized, whereas p197 remains connected to the BB (22, 27). Interestingly, in TAC40-depleted cells ΔC1383-HA was detected in only 20% of all flagella. Thus, most of ΔC1383-HA gets delocalized upon TAC40 depletion suggesting that it binds either to TAC40 itself or to a TAC40-dependent TAC subunit such as TAC42, TAC60 and TAC65. In summary, these results indicate that the N-terminal 365 aa of p197 are sufficient to anchor the protein to the mitochondrial OM. Moreover, since p197 is not affected in the TAC RNAi cell line we can exclude that lateral interaction with full length p197 is responsible for localization of ΔC1383-HA to the TAC region.

**Figure 3.**
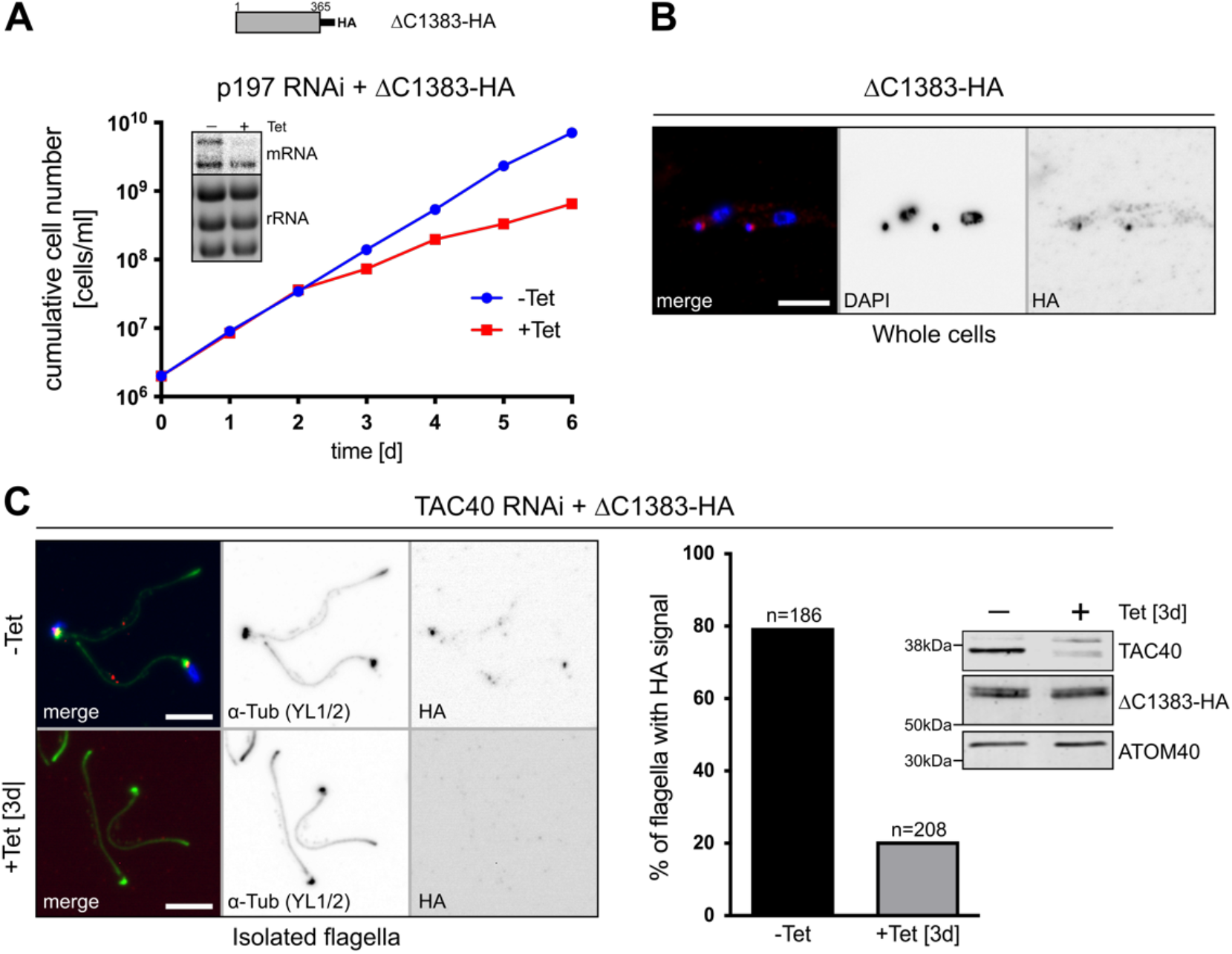
The N-terminus of p197 connects to the OM. **(A)** Growth curve of the p197 3’UTR RNAi cell line expressing an RNAi resistant in-situ C-terminally HA tagged p197 version lacking the 1383 C-terminal aa, termed ΔC1383-HA, schematically depicted at the top. Inset, Northern blot showing ablation of p197 mRNA, ethidiumbromide-stained rRNAs serve as loading control. **(B)** IF analysis of cells expressing ΔC1383-HA stained with DAPI (blue) and for HA (red) as indicated. Scale bar 5μm. **(C)** Left panels, IF analysis of isolated flagella from uninduced and 3 days induced TAC40 RNAi cell line expressing ΔC1383-HA stained with DAPI (blue), the α-tubulin antibody YL1/2 (green) and α-HA (red) as indicated. The contrast of bottom right panel has been enhanced to show that no low insntensity signals are present. Scale bar 5μm. Right panel, quantification of microscopic imaging results. Percentage of flagella with an HA signal was determined with two independent experiments. Number of counted flagella are indicated on top of the bars. Inset, immunoblot confirming ablation of TAC40 upon and constitutive expression of the ΔC1383-HA. ATOM40 serves as a loading control.

### The N-terminus of p197 connects to the peripheral OM protein TAC65

Sequential digitonin fractionations show that p197-HA is almost exclusively recovered in the pellet 2 (P2) fraction indicating that full length p197 is insoluble under these conditions (Fig. 4A). TAC40 behaves similarly, although approximately 50% of the protein is released into the supernatant 2 (SN2). In contrast to full length p197 and TAC40, ΔC1383-HA which is not functional but still can be anchored to the OM (Fig. 3), is completely soluble in 1% digitonin. It behaves like the β-barrel protein ATOM40, a subunit of the mitochondrial OM protein translocase (40), which serves as a control for a digitonin-soluble complex.

**Figure 4.**
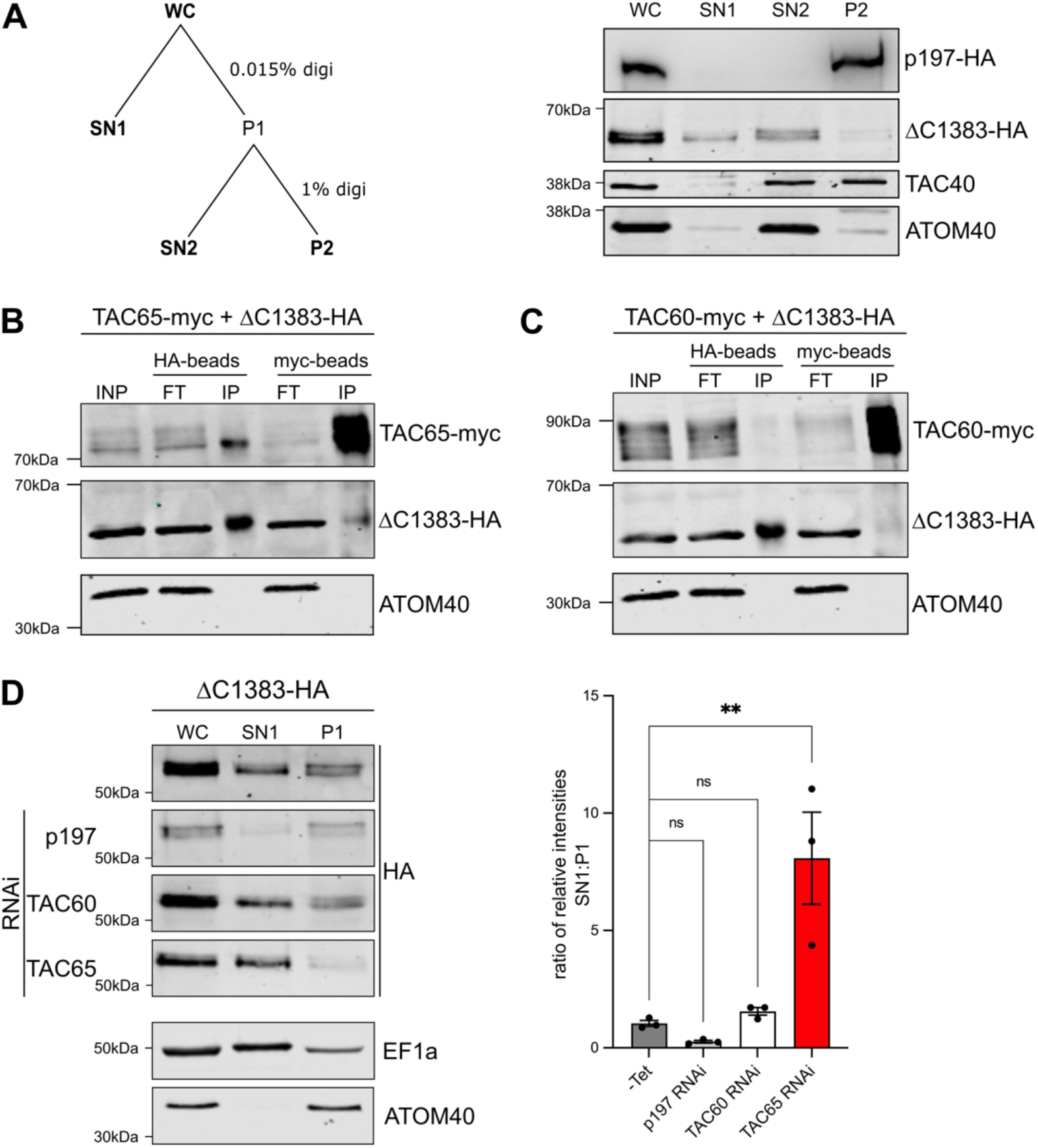
The N-terminus of p197 connects with TAC65. **(A)** Left panel, outline of the digitonin fractionation protocol analyzed by the immunoblot on the right. Right panel, equal cell equivalents of whole cell extract (WC), SN1, SN2 and P2 fractions of cell lines expressing p197-HA and ΔC1383-HA respectively were separated by SDS-PAGE (4% for p197-HA and 12% acrylamide all others) and analyzed by immunoblots. TAC40 and ATOM40 serve as a TAC and mitochondrial marker, respectively. **(B** and **C)** SN2 fractions from cells co-expressing ΔC1383-HA and either TAC65-myc (B) or TAC60-myc (C) were subjected to co-immunoprecipitation using either the anti HA- or anti myc-beads as indicated. 5% of input (INP) and flow through (FT) as well as 100% of the final eluate (IP) were analyzed by immunoblots probed with anti-tag antibodies. ATOM40 serves as negative control for the IP. **(D)** Left panel, immunoblots containing equal cell equivalents of WC, SN1 and P1 fractions from the two days induced p197, TAC60 and TAC65 RNAi cell lines co-expressing ΔC1383-HA were probed for the HA-tagged p197 version and for the presence of the cytosolic and mitochondrial markers EF1a and ATOM40, respectively. Right panel, quantification of the triplicate fractionations shown on the left. The mean of ratio between the signals of SN1 and P1 fractions are indicated. An ordinary one-way ANOVA test with Bonferroni correction has been done. Error bars, standard error of the mean.

To look for putative interaction partners of ΔC1383-HA we produced two cell lines co-expressing ΔC1383-HA together with either C-terminally myc-tagged TAC65 or C-terminally myc-tagged TAC60. Reciprocal Co-IPs demonstrate a stable interaction of TAC65-myc and ΔC1383-HA (Fig. 4B). In contrast, neither TAC60 nor ΔC1383-HA pull down each other (Fig. 4C). These results suggest that binding of ΔC1383-HA to the OM is mediated by TAC65.

The hierarchical assembly model of the TAC (27) proposes that ablation of p197 results in delocalization of all downstream TAC subunits. However, putative assembly intermediates consisting of OM TAC subunits that get dispersed in the OM can still form. Evidence for such putative detergent-soluble assembly subcomplexes containing either TAC40, TAC42 and TAC60 or pATOM36 and TAC65 has been reported (22).

When cells are subjected to subcellular fractionation using a low concentration of digitonin, these OM TAC subunits (together with putative interaction partners) are found in the mitochondria-enriched pellet in contrast to cytosolic proteins that are found in the supernatant. To further corroborate the interaction of ΔC1383-HA with specific OM TAC subunits we biochemically fractionated several cell lines which express ΔC1383-HA and are capable of inducible RNAi targeting p197, TAC60 or TAC65. In uninduced cells substantial amounts of ΔC1383-HA are found in the pellet fraction confirming interaction with OM TAC subunits (Fig. 4D, top panel). Similar results are observed upon downregulation of p197. Ablation of TAC60 leads to the release of small amounts to the soluble supernatant fraction. Strikingly, about two-thirds of the protein are detected in the soluble fraction in the absence of TAC65 which is consistent with the previous conclusion that TAC65 mediates the interaction of ΔC1383-HA with the OM.

In summary, these results strongly suggest that the N-terminus of p197 is orientated towards the OM where it is anchored via the cytosolic and peripherally OM associated TAC subunit TAC65. Furthermore, this proposed interaction does not require more than the N-terminal 365 aa of p197.

### The repeat domain of p197 is essential for function

We investigated whether a shorter version of p197 lacking the entire repeat domain can compensate for the loss of full length p197. Therefore, we stably transfected the existing p197 3’ UTR RNAi host cell line with a construct for inducible expression of an RNAi-resistant variant termed myc-Δrep-HA that lacks the repeat domain but has intact N- and C-termini. The resulting cell line showed a slow growth phenotype 3 days after RNAi induction (*SI Appendix*, Fig. S2A) indicating that the TAC65-binding and BB-binding domains are not sufficient for TAC function but that the central repeat domain p197 is also necessary. IF analyses of isolated cytoskeletons after 3 days of RNAi induction indicate that non-functional myc-Δrep-HA still colocalizes to the BB region, even in the absence of kDNA and the TAC (*SI Appendix*, Fig. S2B). This is expected since the myc-Δrep-HA variant contains the BB-interacting domain and therefore can still bind to the BB.

The simplest explanation for these results is that myc-Δrep-HA is too short to bridge the distance between the BB and the OM.

### p197 bridges the distance between the BB and the OM

The results shown up to now, which are based N- or C-terminally truncated variants of the proteins, strongly suggest that the N-terminus of p197 points towards the OM whereas its C-terminus binds to the BB. However, is this also the case for the complete full length p197? To find out we produced a cell line that exclusively expresses a p197 variant containing both an N-terminal myc-tag and a C-terminal HA-tag (*SI Appendix*, Fig. S3). Figure 5A demonstrates that the resulting cell line, termed myc-p197-HA grows almost identical to wild-type cells indicating that myc-p197-HA is fully functional. In line with this conventional IF analysis shows that both myc and the HA-tag localize to slightly different but partly overlapping regions close to the kDNA (Fig. 5B).

**Figure 5.**
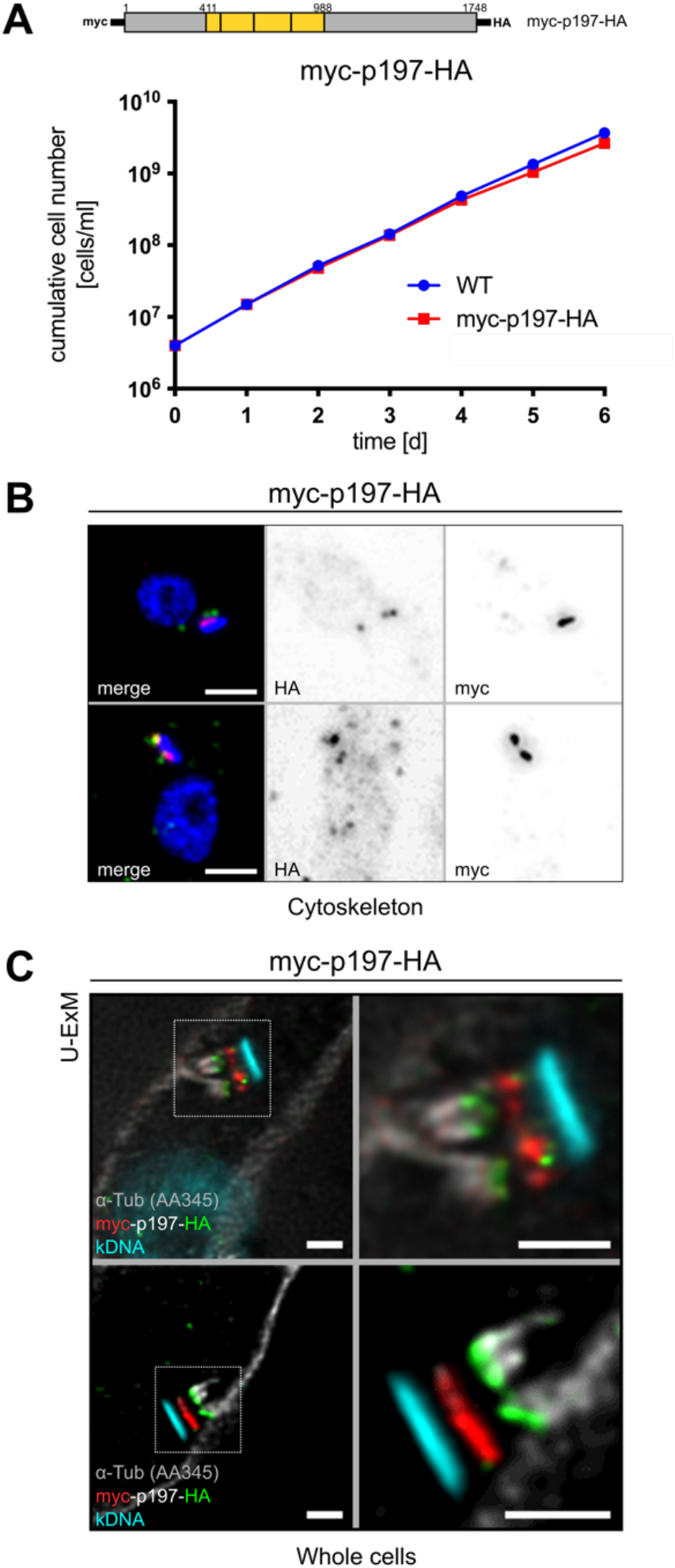
p197 is the exclusion zone filament. **(A)** Growth curve of wildtype cells (blue) and of cells exclusively expressing a N- and C-terminally myc- and HA-tagged p197 version (red), termed myc-p197-HA, schematically depicted at the top. **(B)** Top panels, IF analysis of cytoskeletons isolated from the cell line shown in (A) stained with DAPI (blue), for HA (red) and for myc (green) as indicated. Scale bar 5μm. **(C)** Left panels, U-ExM of myc-p197-HA expressing cells stained with DAPI (magenta), the recombinant α-tubulin antibody AA345 (grey), for HA (green) and for myc (red) as indicated. Right panels, inset of left apanels. Scale bar 2 μm.

To more precisely localize the two tags, we used U-ExM as shown before (Fig. 1). The cells were analyzed with the α-tubulin antibody AA345 to visualize the mature and the pro-BB, with 4’,6-diamidino-2-phenylindole (DAPI) to stain the kDNA disc and with anti-myc and anti-HA antibodies, to localize the N- and C-termini of p197, respectively. The results show that the myc-tagged N-termini (red) localize to a zone in proximity of the mitochondrial OM whereas the HA-tagged C-termini (green) of the same molecules localize to the bottom of mature and pro-BBs (Fig. 5C). The distance that is bridged by p197 can be calculated with the expansion factor (4.6-fold) taken into account. A value of 170 nm after measuring representative images is in line with previously published transmission electron microscopy (TEM) generated images (15). The same holds true for the distance from the kDNA disc to the N-terminus of p197. It roughly corresponds to the kDNA-OM distance. These results are in agreement with the analyses of the truncated p197 variants. They show that the EZFs of the TAC are formed by p197 which bridges the gap from the BBs to the OM in a defined orientation: the N-terminus facing the OM and the C-terminus interacting with the BB region.

## Discussion

In the present study we show that the EZFs as proposed previously (15) are made of filaments consisting likely of single p197 molecules spanning the region from BBs to the periphery of the mitochondrial OM (see model Fig. 6). We furthermore show that assembly of these filaments starts very early in the cell cycle, immediately after the formation of a new pro-BB before the new flagellum starts to grow.

**Figure 6.**
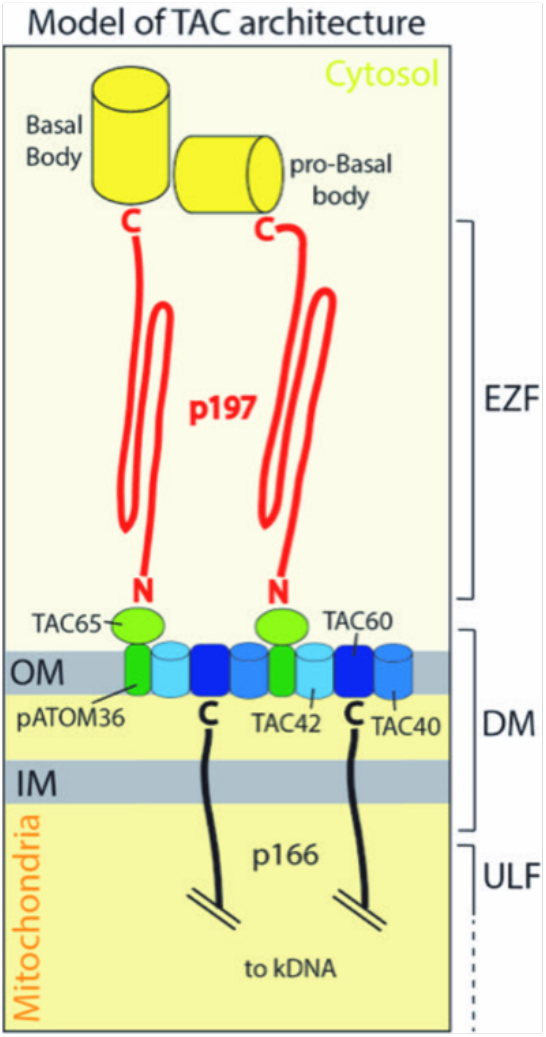
Model depicting the architecture of the EZF and the DM region of the trypanosomal TAC. p197 likely is the TAC subunit most proximal to the BB. One p197 molecule (red) each that connects either the BB or the pro-BB to the OM is shown. The C-terminus of p197 interacts with the BB or the pro-BB and its N-terminus binds to the cytosolic membrane peripheral TAC65. p197 is predicted to have a high α-helical content which likely forms coiled coil domains containing multiple α-helices. TAC65 and pATOM36 form a subcomplex in OM that is indicated in green. TAC60, TAC42 and TAC40, the latter two of which are β-barrel proteins, from a second OM subcomplex that is depicted in blue. p166 interacts with TAC60 via its C-terminus. It has a single transmembrane domain and likely is the only TAC subunit integral to the IM. Thus, the TAC65/pATOM36 and the TAC60/TAC42/TAC40 subcomplexes form a platform in the OM that integrates the EZFs with the intramitochondrial ULFs. How the ULFs connect to the kDNA is not shown.

p197 can be divided into three domains consisting of the N-terminal, the C-terminal and the central region (Fig. *SI Appendix*, S4) all of which have distinct functions and are required for an intact TAC. TAC65 is a soluble protein that is peripherally associated with the mitochondrial OM. It has originally been identified in a pull-down experiment of tagged pATOM36 that also recovered p197 (22). The OM protein pATOM36 is an essential TAC subunit and additionally required for the biogenesis of a subset of mitochondrial OM proteins (23, 25). Reciprocal pull-down analyses using tagged TAC65 as a bait recovered pATOM36 suggesting that TAC65 and pATOM36 form a complex that likely also contains p197. Here we show that TAC65 anchors p197 to the differentiated OM by binding to the N-terminal region of p197 and to the OM TAC subunit pATOM36 (see model Fig. 6).

U-ExM indicates that the p197 filaments originate from or in the vicinity of the ring defined by the 9 triplet microtubules at the bottom of the mature and the pro-BB suggesting that p197 is the TAC subunit most proximal to the BB. The orientations of the old and the new mature and pro-BB are dynamic and sometimes perpendicular to each other. Thus, there is an anti-clockwise rotation of the new mature BB around the old flagellum (41). p197 is attached to the mature and pro-BB throughout the cell cycle indicating that it must be a flexible molecule that can be bent. To which BB component the C-terminal region of p197 remains to be determined.

The central region of p197 consisting of the 175 aa repeats make up a large portion of the protein. However, it is difficult to determine the exact repeat number since near identical tandem repeats often lead to genome assembly errors (42) and therefore the correct size and molecular mass of p197 are unclear. The following lines of evidence suggest that p197 is much larger than 197 kDa. Recent sequencing of the genome of the *T. b. brucei* EATRO1125 strain revealed that p197 has at least 26 repeats of 175 aa (43), indicating that the molecular weight of p197 might be as large as 660 kDa. That would make it to one of the largest trypanosomal proteins known. This might be similar in the *T. brucei* 427 strain used in the present study as evidenced by SDS PAGE shown in Figure S1C which shows that the molecular mass of p197 is much higher than the highest molecular weight marker of 260 kDa.

The Phyre2 algorithm predicts that the repeat domain of p197 has an α-helical content of 96% (see *SI Appendix*, Fig. S5A). Considering a molecular weight of 660 kDa and assuming p197 consists mainly of α-helices it could span a maximum distance of 853 nm (3.6 aa and 0.54 nm per helix turn, respectively). The actual distance between the BB and the OM that has to be spanned by p197 is difficult to estimate since in agreement with the predicted flexibility of the protein it is quite variable. Measurements using a variety of previously published TEM pictures suggest it to be approximately 150 nm (15). Thus, the predicted length of p197 would be sufficient to bridge this distance more than 5-times. However, it is likely that the central domain of p197 does not simply adopt an α-helical structure because the diameter of the EZF filaments has been measured to be between 5 to 10 nm (15) which is larger than the 1.2 nm diameter of an average α-helix. Thus, we expect the central domain of p197 to adopt a higher order structure possibly including coiled coil domains. Removing the repeat domain from p197, as has been done in the experiment shown in Fig. S2, shortens p197, assuming its true molecular weight is 660 kDa, by a factor of more than three. This suggests that the resulting protein is simply too short to span the required distance from the BB to the OM.

Our work also illustrates the power of U-ExM which was done under denaturing conditions but in the absence of proteases. In a cell line which exclusively expresses double tagged p197, the myc- and the HA-tag localize to OM and BB regions of the TAC, respectively. The simplest explanation for this result is that we succeeded to visualize single p197 molecules that are arranged in a parallel bundle. This suggests that the expansion polymer in U-ExM is very uniformly distributed allowing for high precision expansion. The linear expansion factor in Fig. 5C is 4.6 which results in a BB-OM distance of 730-780 nm. This is still short enough to be bridged by p197 in an α-helical formation. We therefore suggest that in our U-ExM the p197 molecules get stretched out and that at least some of the postulated coiled coil domains are resolved. The alternative that the protein chain is ripped apart is unlikely since the covalent bonds of the protein backbone are much stronger than the non-covalent interactions forming coiled coil domains.

The kDNA and its connection to the flagellum via the TAC is a defining feature of the *Kinetoplastidae*. Thus, orthologues of p197 are found in essentially all kinetoplastids.

A multiple sequence alignment of p197 with a group of diverse Kinetoplastids comprising *T. brucei, Trypanosoma cruzi, Leishmania major* and *Paratrypanosoma confusum* revealed that the N-terminal 150 aa as well the 60 aa of the very C-terminus are more conserved than the central region of the protein (*SI Appendix*, Fig. S6). All p197 orthologues are of high predicted molecular mass and have an overall high α-helical content (*SI Appendix*, Fig. S5B). However, while the *T. brucei* p197 contains nearly identical repeats no such repeats are found in the central region of p197 orthologues of other kinetoplastids. The N-and C-termini of p197 are involved in protein protein interaction to the OM and the BB respectively. This may explain why these regions are under a stronger selection pressure and more conserved than the diverged central part of p197, which likely just needs to provide an α-helical flexible linker of sufficient length.

## Material and methods

### Transgenic cell lines

All procyclic cell lines are derivatives of *T. brucei* 427 grown at 27°C in SDM-79 supplemented with 5% (v/v) fetal calf serum (FCS). To generate a single marker inducible cell line, 427 cells were transfected with a modified pSmOx plasmid (44) where the puromycin resistance gene had been replaced with phleomycin. The plasmid encodes both the T7 RNA polymerase (RNAP) and the Tet-repressor.

For Tet-inducible RNAi of p197 (Tb927.10.15750) cells were stably transfected with a NotI-linearized plasmid containing stem-loop sequences covering the 3’ UTR of p197. RNAi plasmids targeting the ORFs of TAC65, TAC60 or TAC40 have been described previously (22, 23).

*In situ* C-terminal HA-tagging of p197 was done by stable transfection of PCR products using plasmids of the pMOtag series as templates (45) with primers defining the sites of homologous recombination leading to expression of tagged full length p197 or versions thereof lacking 334aa or 1383 aa of its C-tail. *In situ* N-terminal myc-tagging of p197 was also done using PCR products with primers defining the flanking region of the start codon of p197. The plasmid for amplification was a derivate of the pN-PURO-PTP (46) and was modified to have a triple myc tag instead of the PTP tag.

Plasmids for tetracycline-inducible expression of ΔC1383-HA, ΔN1228-HA, TAC65-myc and TAC60-myc were generated with a modified pLew100 expression vector (47) containing a puromycin or a blasticidin resistance cassette. For p197 truncations, ORFs that either lacked the first 1228 or the last 1383 aa were amplified from genomic DNA and integrated into the vector allowing C-terminal triple HA tagging. For TAC65 and TAC60 the whole ORF lacking the stop codon was amplified from genomic DNA and integrated into the vector allowing for C-terminal triple myc tagging.

To ensure functionality of double-tagged p197, a plasmid for the single knockout of p197 was generated by modifying a pMOtag43M vector (45). The 5’ flanking sequence of p197 (500 nucleotides before start codon) and the 3’ flanking sequence (495 nucleotides after stop codon) were amplified and ligated into XhoI/NheI and BamHI/SacI restriction sites of the vector thereby flanking the hygromycin resistance gene. Because the 3’ flank of p197 has an internal XhoI restriction site (starting at nucleotide 226 after stop codon) the resistance cassette could be released with XhoI before electroporation of the single or double tagged p197 cell line. For an overview of p197 variant used in this study see *SI Appendix*, Figure S6.

### Immunofluorescence microscopy

1×10^6^ cells were harvested, washed, resuspended in phosphate buffered saline (PBS) and allowed to settle on cover slides for 10 min. For whole cell IF analysis parasites were fixed using 4% paraformaldehyde (PFA) and permeabilized with 0.2% Triton X-100. For cytoskeletons cells were first permeabilized with 0.05% Triton X-100 for 10s, washed in PBS and then fixed with 4% PFA. Flagella were isolated as described (48). In brief, cells were harvested by centrifugation after EDTA (100nM) was added to the culture, directly resuspended in extraction buffer (10mM NaH_2_PO_4_, 150mM NaCl, 1mM MgCl_2_, pH 7.2) containing 0.5% Triton X-100 and incubated for 10 min on ice. After centrifugation pellets were incubated on ice for 45min with extraction buffer containing 1mM CaCl_2_. Extracted flagella could then be pelleted for 10min at 10’000g, resuspended in PBS, settled on a coverslip for 10min and fixed with 4% PFA.

Staining with antibodies has been done in a wet chamber, primary antibodies were diluted in PBS containing 2% bovine serum albumin (BSA), secondary fluorescent antibodies in PBS only. After incubation in cold methanol, dried coverslips were mounted onto Vectashield containing DAPI (Vectorlabs).

Pictures were acquired using a DMI6000B microscope equipped with a DFC360 FX monochrome camera and LAS X software (Leica Microsystems). Images were further analyzed using Fiji software and the FigureJ Plugin.

### Ultrastructure expansion microscopy (U-ExM)

80μl containing two million cells were settled for 20min at room temperature on poly-D-lysine functionalized coverslips (12mm, Menzel-Glaser). Coverslips were transferred into a 24-well plate filled with a FAB solution (0.7% Formaldehyde (FA, 36.5-38%, Sigma) with 0.15% or 1% acrylamide (AA, 40%, Sigma) in PBS) and incubated overnight at 37°C. Cells were then prepared for gelation by carefully putting coverslips from the FAB solution into a 35μl drop of monomer solution (AK Scientific 7446-81-3) 10% (wt/wt) AA, 0.1% (wt/wt) N,N’-methylenebisacrylamide (BIS, Sigma) in PBS supplemented with 0.5% ammoniumpersulfate and 0.5% tetramethylethylendiamine (TEMED) on parafilm in a pre-cooled humid chamber. Gelation was done for seven minutes on ice, and subsequently samples were incubated at 37°C in the dark for one hour. Coverslips with gels were then transferred into a six well plate filled with denaturation buffer (200mM SDS, 200mM NaCl, and 50mM Tris, pH 9 in ultrapure water) for 15min at room temperature. Gels were then detached from the coverslips with tweezers and moved into a 1.5ml Eppendorf centrifuge tube filled with denaturation buffer and incubated at 95°C for one hour and 30min. After denaturation, gels were placed in deionized water for the first round of expansions. Water was exchanged at least twice every 30min at room temperature and incubated overnight in deionized water. On the next day, gels were washed two times for 30min in PBS and subsequently gently agitated on a shaker with the primary antibody diluted as follows: guinea pig α-tubulin antibody detecting (AA345, Geneva Antibody Facility) 1:500; anti-HA rabbit (H6908, Sigma) 1:500, anti-myc mouse (M4439, Sigma) 1:500. All antibodies were diluted in 2% PBS/BSA and gels were incubated for 3 hours at 37°C. Gels were then washed in PBST three times for 10min while gently shaking and subsequently incubated with secondary antibodies Alexa Fluor 488 goat-anti-rabbit IgG (H+L) (Invitrogen), Alexa Fluor 594 goat-anti-mouse IgG (H+L) (Molecular probes), Alexa Fluor 647 goat-anti-rat IgG (H+L) (Life technologies),anti-guinea pig 647 (Abcam 150187) all diluted 1:1’000, and DAPI (Molecular Probes) diluted 1:5’000. All antibodies were diluted in PBS containing 2% BSA and gels were incubated for approximately 3 hours at 37 °C. Gels were then washed in PBST three times for 10 minutes while gently shaking and finally placed in beakers filled with deionized water for expansion. Water was exchanged at least twice every 30 minutes before gels were incubated in deionized water overnight.

### Mounting and image acquisition

After the final expansion gel size was measured with a caliper to calculate the fold expansion. The gel was then cut with a razor blade in pieces that fit in a 36mm metallic chamber for imaging. The gel piece was then mounted on a 24mm round poly-D-lysine functionalized coverslip, already inserted in the metallic chamber, and gently pressed with a brush to ensure adherence of the gel to the coverslip. Confocal microscopy was performed on a Leica TCS SP8 using a 63 × 1.4 NA oil objective with the following parameters: z step size at 0.3μm interval with a pixel size of 35nm. LAS X software (Leica Microsystems), Huygens Professionals and ImageJ were used to analyse the images.

### Calculation of expansion from U-ExM

The expansion factor was determined by measuring the expanded lengths of the cell, the kDNA and the diameter of the nucleus and comparing these values to the ones in non-expanded samples. kDNA was measured by scoring the maximum length observed in each cell. The diameter of the nucleus was defined as the widest diameter observed in each cell. The length of the cell structure was determined by measuring their distance from the anterior to the posterior of each cell. Twenty cells each were analyzed. The reached linear expansion factors were between 4.2 and 4.6.

### Cell fractionation

Two-step digitonin extractions were performed to analyze the subcellular localization of proteins. 1×10^7^ cells were harvested, washed and resuspended in SoTE buffer (20mM Tris HCl pH 7.5, 0.6M sorbitol, 2mM EDTA). An equal volume of SoTE containing 0.03% (w/v) digitonin was added for selective cell membrane permeabilization. After incubation on ice for 10min the sample was centrifuged for 5min at 6’800g. The supernatant (cytosolic fraction, SN1) was stored on ice and the pellet was then subjected to 1% digitonin extraction, left on ice for 15min and centrifuges at 21’000g for another 15min. Equal cell equivalents of the resulting supernatant (soluble organellar proteins, SN2), the pellet (insoluble proteins, P2), SN1 fraction and whole cell fractions (WC) were analyzed using SDS-PAGE.

### Immunoprecipitations

For co-immunoprecipitation 2×10^8^ cells were used to generate a mitochondria-enriched pellet using 0.015% of digitonin. The pellet was then solubilized by incubation in lysis buffer (20 mM Tris HCl pH 7.4, 100 mM NaCl, 0.1 mM EDTA, 10% glycerol) containing 1% digitonin and a protease inhibitor cocktail (Roche Complete, EDTA free). After 15 min on ice, the sample was centrifuged for 15min at 21’000g. Half of the supernatant was incubated with anti-c myc beads (Sigma) or anti-HA beads (Roche) for 1 hour at 4°C. Afterwards, the flow-through was collected, beads were washed 3 times with lysis buffer containing 0.1% digitonin and proteins were eluted by boiling the beads in SDS-PAGE sample loading buffer lacking β-mercaptoethanol. Fractions of the input, the flow-through and the entire eluted sample were subjected to immunoblot analysis.

### RNA extraction and Northern blotting

Total RNA was extracted using acid guanidinium thiocyanate-phenol-chloroform according to (49) and separated on a 1% agarose gel. Northern blot probes for the corresponding RNAi targeting fragment were radioactively labeled using the Prime-a-Gene labelling system (Promega).

### Antibodies

The following antibodies were used in the study. The dilution for immunoblots (IB) and IF is indicated in parentheses. To produce a polyclonal rabbit antiserum against TAC40 (IB: 1:100, IF 1:100), the entire ORF of the protein containing an N-terminal His-tag, was recombinantly expressed in *E. coli*, purified via nickel columns and used to immunize rabbits (Eurogentec). The polyclonal rabbit antiserum against ATOM40 (IB 1:10’000, IF 1:1,000) has previously been described (50). The polyclonal rabbit anti SAS6 antiserum (IF 1:400) was a gift from Prof. Ziyin Li and the rat monoclonal α-tubulin YL1/2 (IB 1 :5’000, IF 1 :1000) was a kind gift from Prof. Keith Gull. The following commercially available monoclonal antibodies were used in this study: monoclonal mouse anti-myc antibody (Invitrogen, 132500; IB 1:2’000, IF 1:50), monoclonal mouse anti-HA antibody (Enzo Life Sciences AG, CO-MMS-101 R-1000; IB 1:5’000, IF 1:1’000), polyclonal rabbit anti HA antiserum (Santa Cruz, Y-11, IF 1:250) and monoclonal mouse anti-EF1a antibody (Merck Millipore, product no. 05– 235; WB 1:10’000).

Secondary antibodies for IB analyses were IRDye 680LT goat anti-mouse, IRDye 800CW goat antirabbit (LI-COR Biosciences; 1:20’000), and HRP-coupled goat anti-mouse (Sigma-Aldrich; 1:5’000). Secondary antibodies for the IF analysis were goat anti-mouse Alexa Fluor 596, goat anti-rat Alexa Fluor 488, goat anti-rabbit Alexa Fluor 488 (all from ThermoFisher Scientific; IF 1:1’000).

### Miscellaneous

Immunoblots- and Northern blots were done as previously described (51).

## SI Appendix: Supplementary Figure S1-S6

**Supplementary Figure S1.**
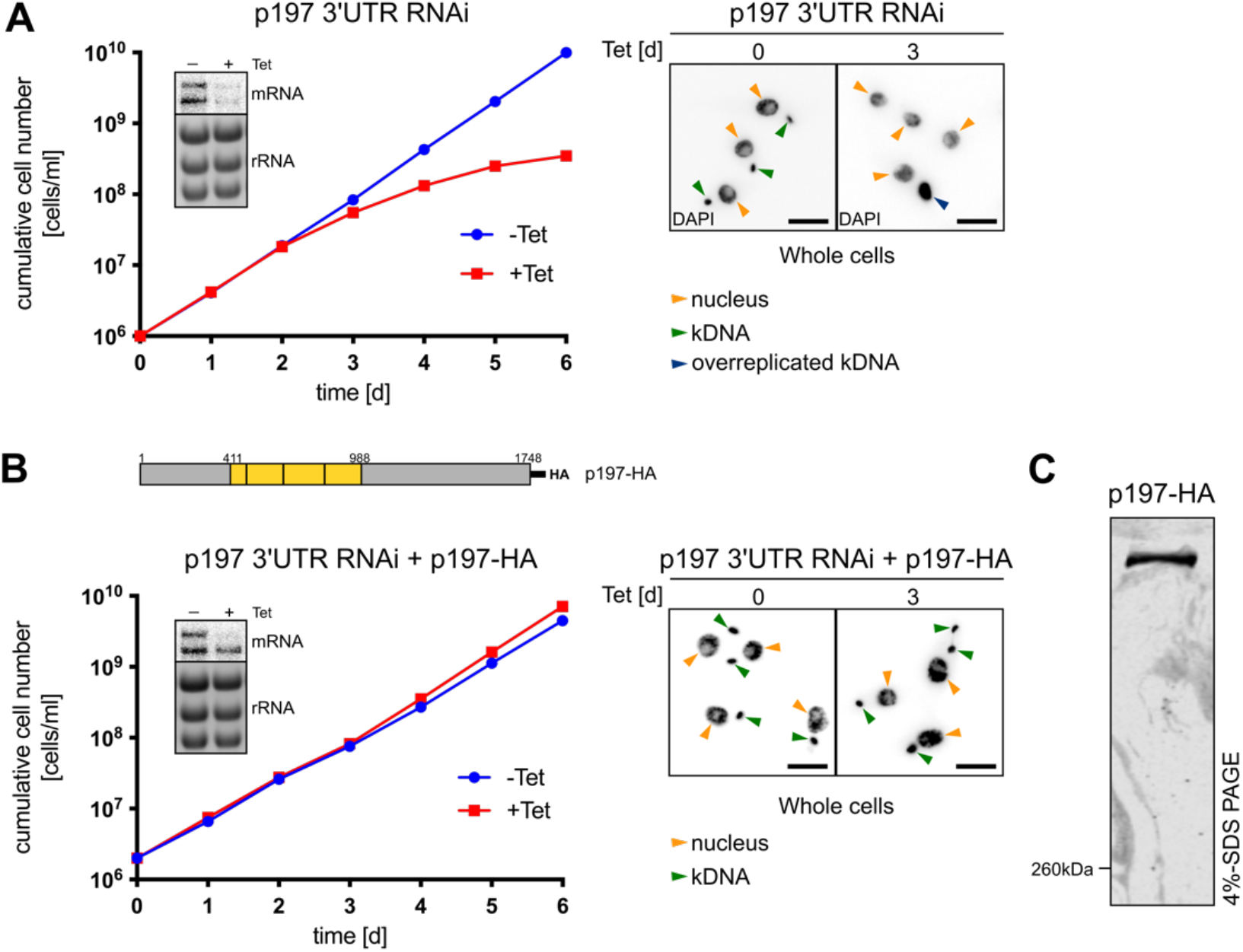
C-terminally HA tagged p197 is functional. **(A)** Left panel, growth curve of the tetracycline (Tet) inducible p197 3’UTR RNAi cell line. Inset, Northern blot probed for the p197 mRNA. Ethidium bromide-stained rRNAs serve as loading control. Right panel, DAPI staining of whole cells of the p197 3’UTR RNAi cell line. Nuclei (orange), normal sized kDNAs (green) and overreplicated kDNAs (blue) are indicated with arrowheads. Scale bar 5μm. **(B)** Growth curve of the p197 3’UTR RNAi cell line expressing an RNAi-resistant in-situ C-terminally HA tagged p197 version schematically depicted at the top. Yellow, 175 aa repeats. Right panel, DAPI staining and scale bar as in (A). **(C)** Immunoblot of HA-tagged p197 from whole cell extracts separated by 4% SDS PAGE.

**Supplementary Figure S2.**
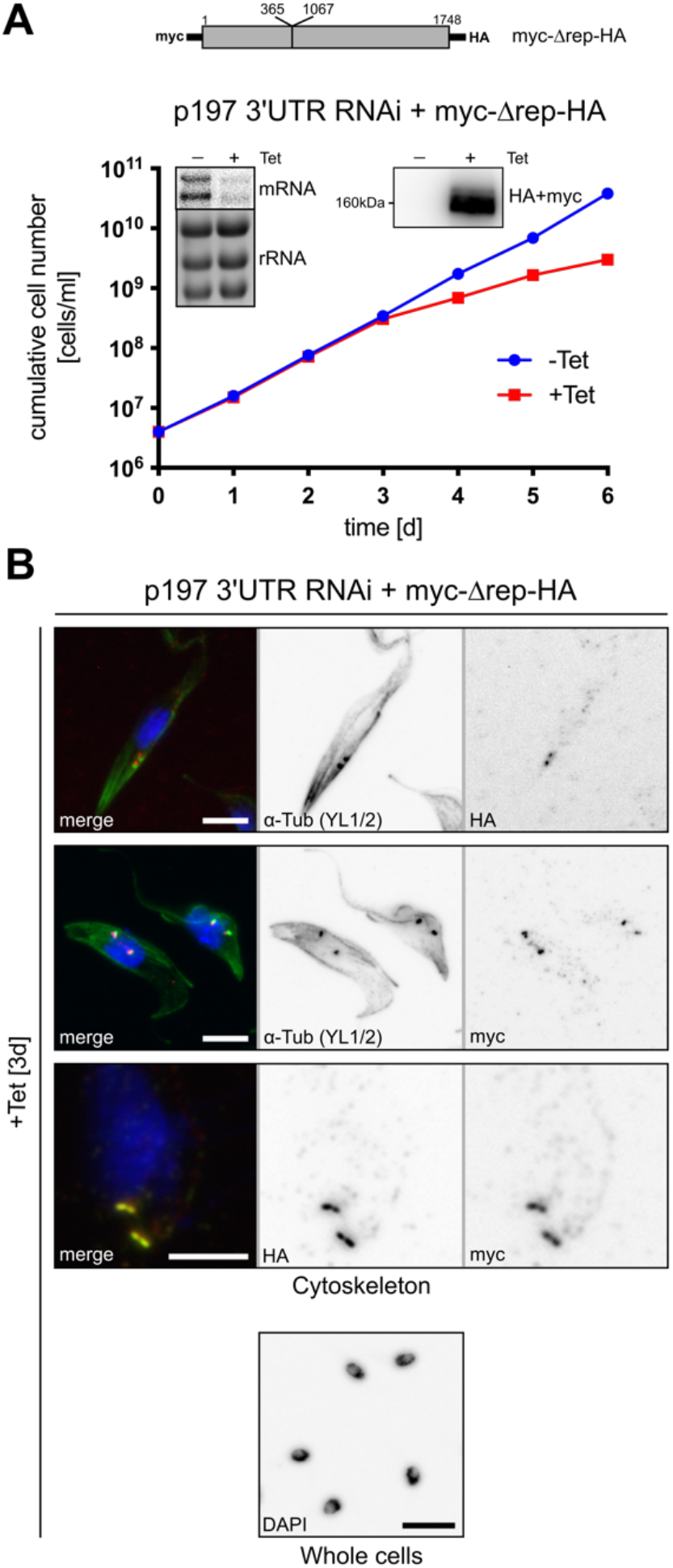
The repeat region is essential for p197 function. **(A)** Growth curve of the p197 3’UTR RNAi cell line expressing an RNAi resistant tet-inducible N- and C-terminally myc- and HA-tagged p197 version lacking the central repeat domain, termed myc-Δrep-HA, schematically depicted at the top. Inset left, Northern blot showing ablation of p197 mRNA, ethidiumbromide-stained rRNAs serve as loading control. Inset right, immunoblot showing inducible expression of myc-Δrep-HA. **(B)** Top panels, IF analysis of cytoskeletons isolated from the cell line shown in (A) stained with DAPI (blue), the α-tubulin antibody YL1/2 (green), for HA (red) and for HA and myc simultaneously (red) as indicated. Bottom panel, IF analysis of whole cells corresponding to the top panels stained with DAPI to illustrate the RNAi-induced kDNA loss. Scale bar 5μm.

**Supplementary Figure S3.**
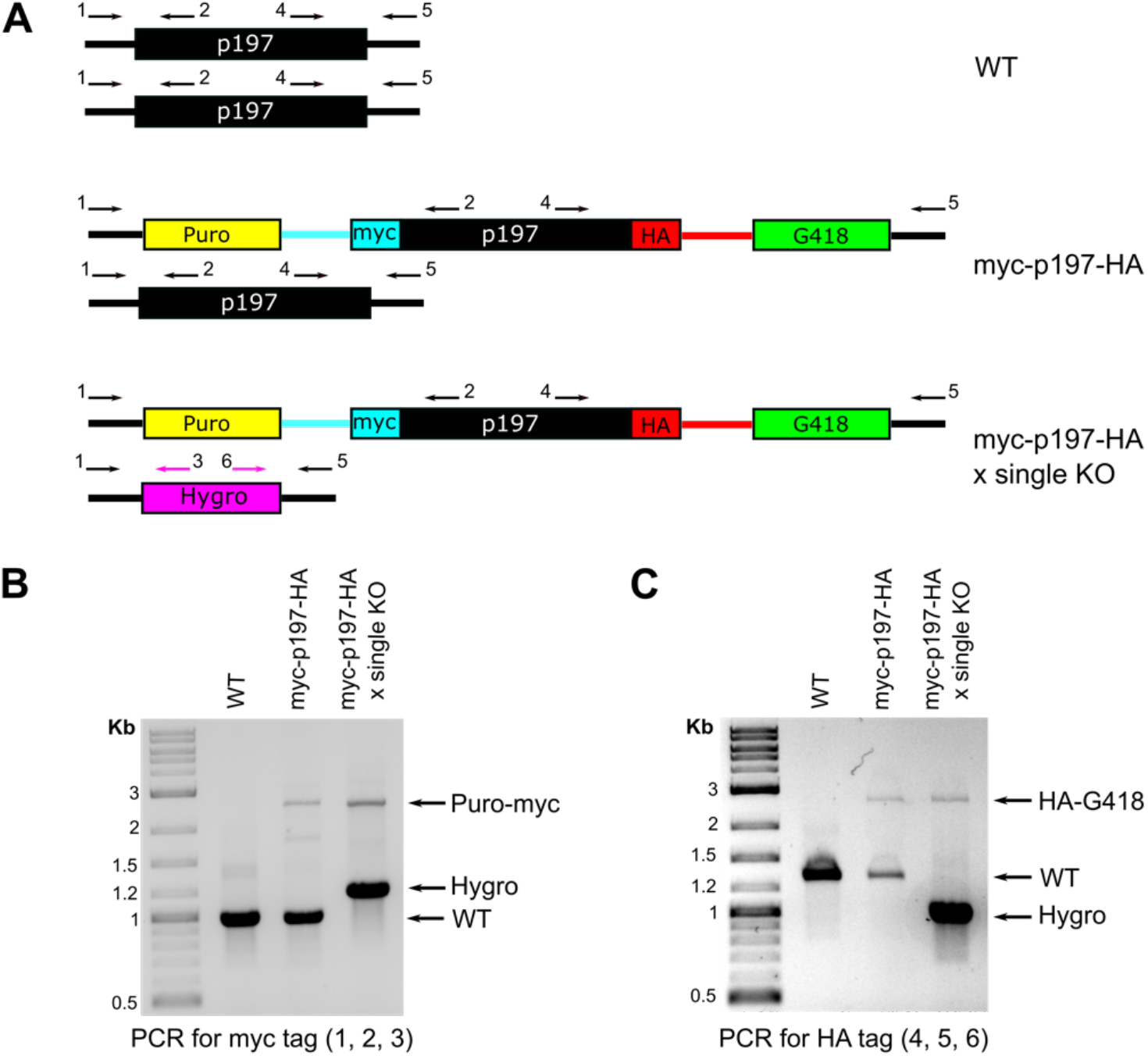
Exclusive expression of myc-p197-HA. **(A)** Allelic situation of either wildtype cells (WT), cells containing double tagged p197 in the background of one wild-type allele of p197 (myc-p197-HA), or cells containing double tagged p197 in the background of a single knockout of p197 (myc-p197-HA x single KO) is schematically depicted (not to scale). Arrows and numbers indicate PCR primers that have been used for verification of the cell lines. **(B** and **C)** Genomic DNA of the three cells lines described in (A) was isolated and used as templates for the PCR reactions. A competition PCR with three primers (indicated with numbers) was used to verify the presence of the tag-containing constructs and the single knock outs of p197 on the genomic level. The presence of the N- and C-terminal tag encoding DNA sequences was verified by two independent PCR reactions. The myc-p197-HA x single KO cell line lacks a PCR product that in size corresponds to the one in WT cells, instead an additional PCR product for hygromycin is detected. In summary this shows successful knockout of the remaining p197 wild type allele.

**Supplementary Figure S4.**
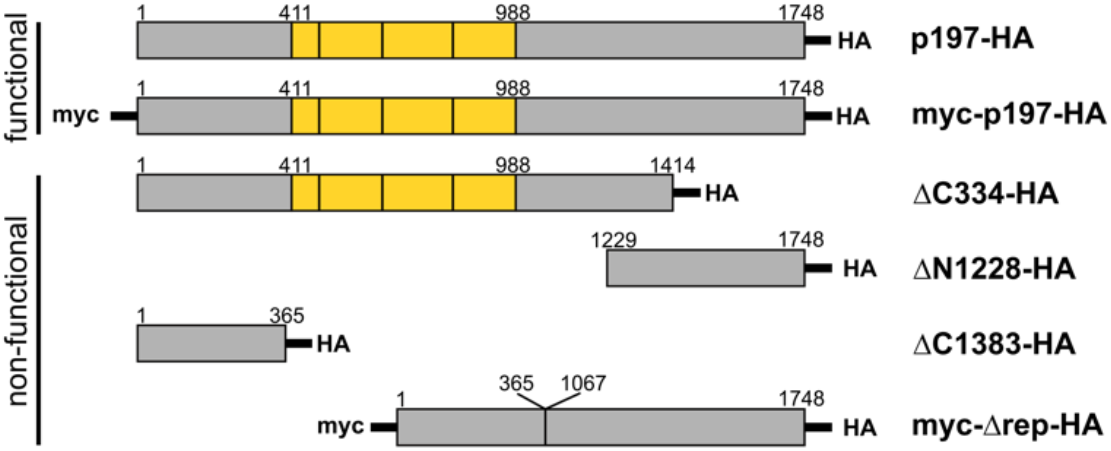
Schematic representations of p197 versions. p197 variants that have been used in this study are shown schematically. Yellow highlights the repeat region. Constructs complementing TAC function are indicated. All p197 variants localized to the TAC region.

**Supplementary Figure S5.**
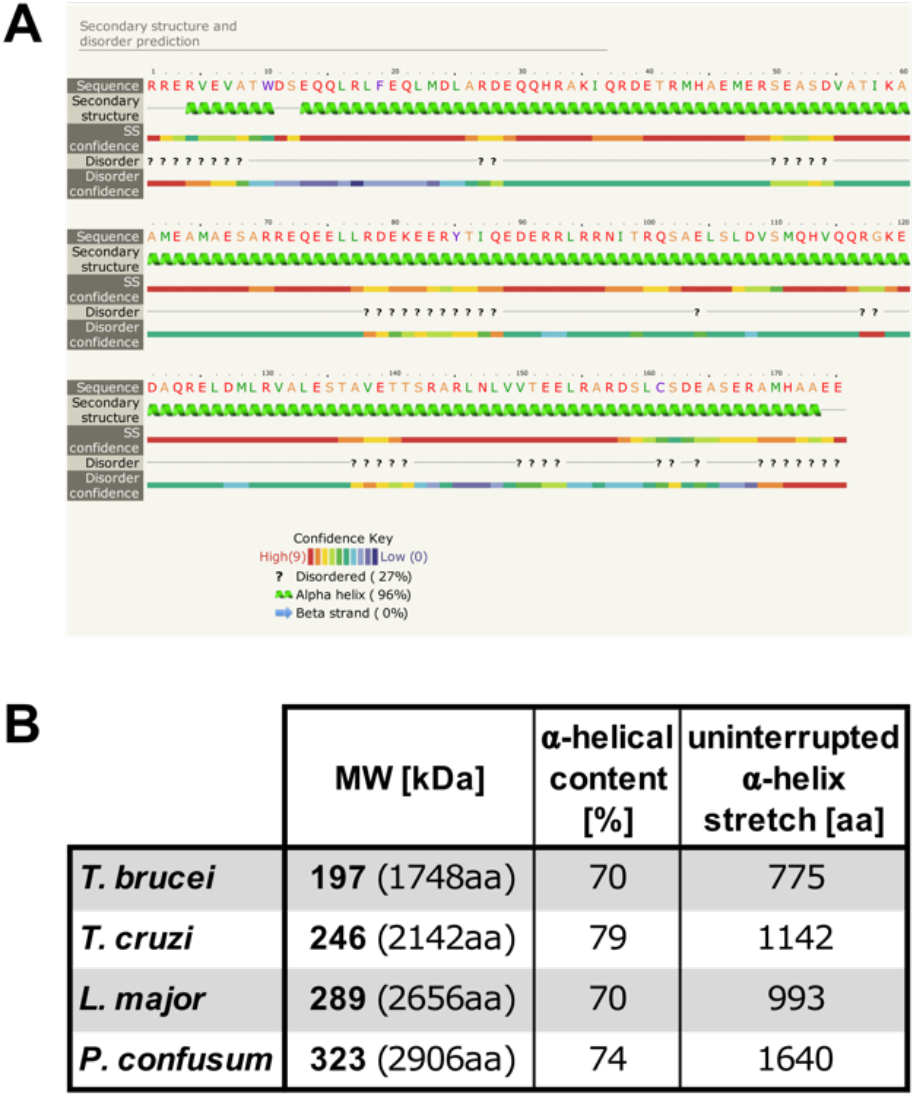
p197 orthologues have a largely α-helical structure. **(A)** Secondary structure of one 175 aa repeat from *T. brucei* p197 has been predicted using the Phyre2 online tool. It has an α-helix content of 96%. **(B)** Comparison of p197 homologues from the indicated organisms. Calculated molecular weights (with length of the proteins in parentheses), α-helical content and length of the longest essentially uninterrupted α-helical stretch are indicated.

**Supplementary Figure S6.**
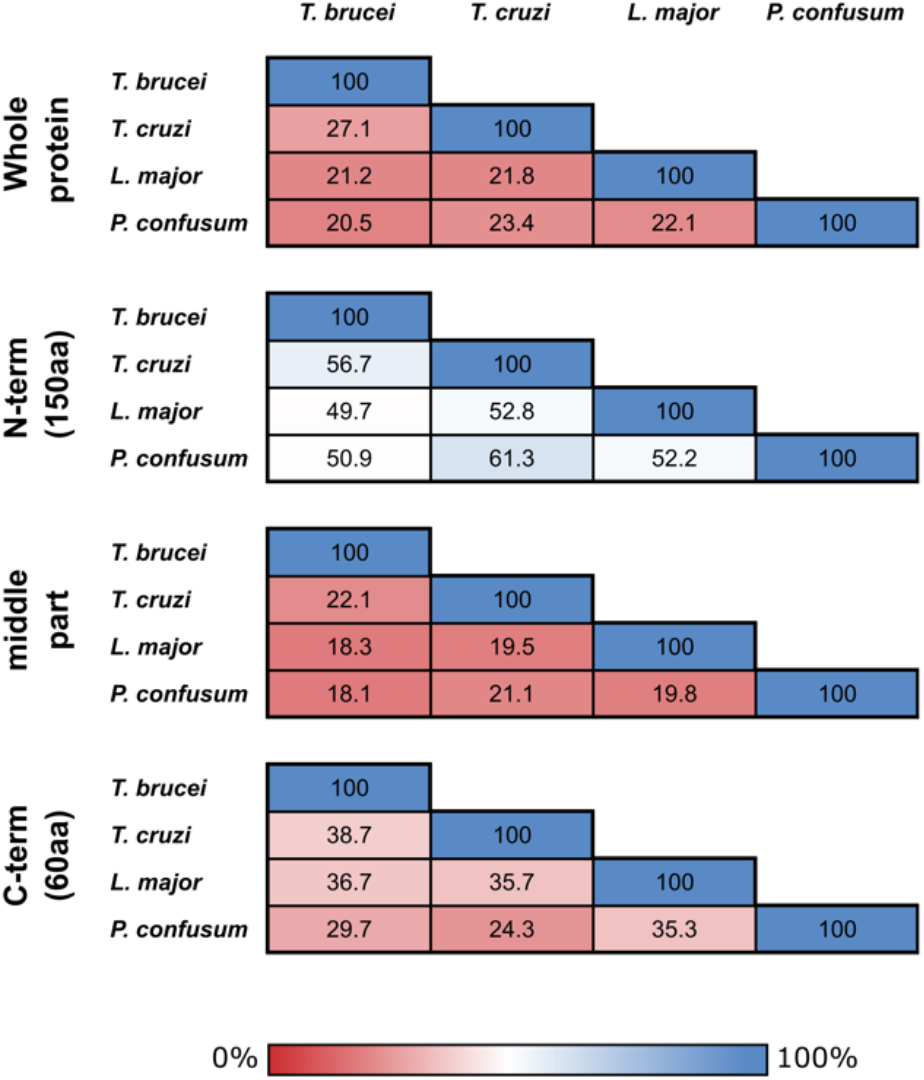
The N- and C-termini of p197 are more conserved than the rest of the protein. Full length p197 orthologous or indicated parts of them from *T. brucei, Trypanosoma cruzi, Leishmania major* and *Paratrypanosoma confusum* were pairwise compared using BLAST. Percentage of identity is indicated.

## Notes

### Competing Interest Statement

The authors have declared no competing interest.

